## Supplemental table 2 for "Identification of Serum Bridging Molecules that Mediate Human Endothelial Cell Invasion by *Candida* species"

**Table S2.** Fungal strains used in this work.

| Organism | Strain | Relevant Genotype | Source/  Reference |
| --- | --- | --- | --- |
| *Candida albicans* | SC5314 | Wild-type | (1) |
| *Candida albicans* | DIC185 | *ura3*Δ::λ*imm434/ura3*Δ::λ*imm434*::*URA3-IRO1*  *arg4*::*hisG*/*arg4*::*hisG*::p*ARG4*  *his1*::*hisG/his1*::*hisG*::p*HIS1* | (2) |
| *Candida albicans* | CAI4-URA | *ura3*Δ*::imm434*/*ura*3Δ*::imm434::URA3* | (3) |
| *Candida albicans* | DSC10 | *efg1*Δ*::hisG*/*efg*1Δ*::hisG*  *ura3*Δ*::* λ*imm434*/*ura*3Δ*::* λ*imm434::URA3-IRO1* | This study |
| *Candida albicans* | *ssa1*Δ/Δ-*als3*Δ/Δ | *ura3*Δ::λ*imm434/ura3*Δ::λ*imm434*  *ssa1*Δ::*FRT/ssa1*Δ::*FRT*  *SSA2/ssa2*Δ::*FRT*  *als3*Δ/*als3*Δ*::NAT1*  *RPS10/rps10::URA3* | (4) |
| *Candida auris* | CAU-09 | Wild-type | (5) |
| *Candida glabrata* | 05-761 | Wild-type | (6) |
| *Candida krusei* | 2333 | Wild-type | Fungus Testing Laboratory at the University of Texas Health Science Center at San Antonio |
| *Candida parapsilosis* | 1184 | Wild-type | Fungus Testing Laboratory at the University of Texas Health Science Center at San Antonio |
| *Candida tropicalis* | DI13-280 | Wild-type | Fungus Testing Laboratory at the University of Texas Health Science Center at San Antonio |
| *Saccharomyces*  *cerevisiae* | AMP188 | *MAT*α *lys-23 ade2 ura2-49* (Σ1278b background) | (7) |
