## Supplemental figures for "Identification of Serum Bridging Molecules that Mediate Human Endothelial Cell Invasion by *Candida* species"

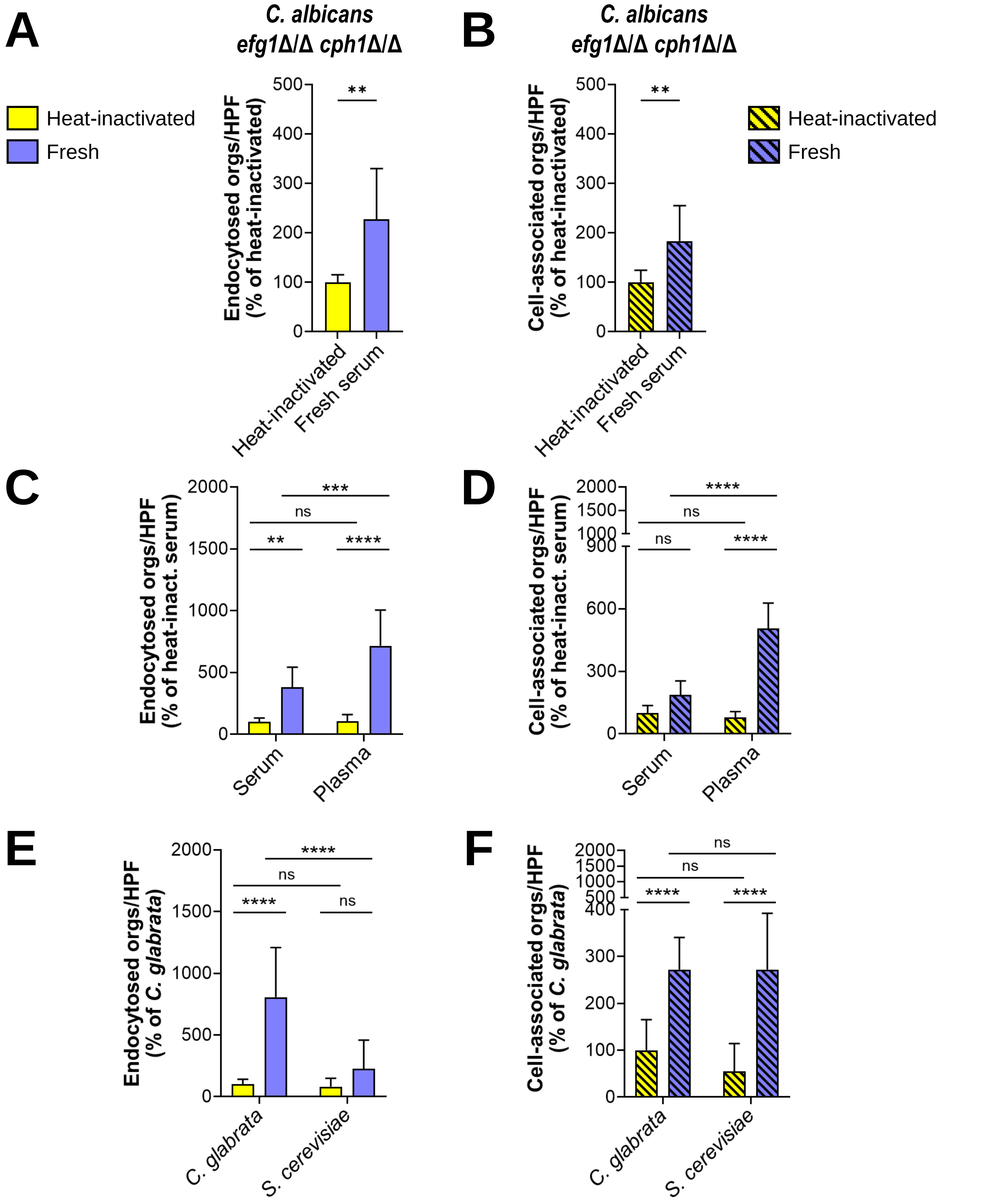


**Fig. S1.** Bridging molecules mediate the endocytosis of live *C. albicans* and *C. glabrata* yeast, but not *S. cerevisiae.* (A and B) Endocytosis (A) and cell-association (B) of live cells of a *C. albicans egf1*Δ/Δ *cph1*Δ/Δ mutant by human umbilical vein endothelial cells. (C and D) Effects of fresh human serum and plasma on the endocytosis (C) and cell-association (D) of live *C. glabrata*. (E and F) Endocytosis (E) and cell-association (F) of live *C. glabrata* and *S. cerevisiae.* Results are the mean ± SD of 3 independent experiments, each performed in triplicate. Orgs/HPF, organisms per high-power field; ns, not significant; **$P < 0.01$, ***$P < 0.001$, ****$P < 0.0001$.

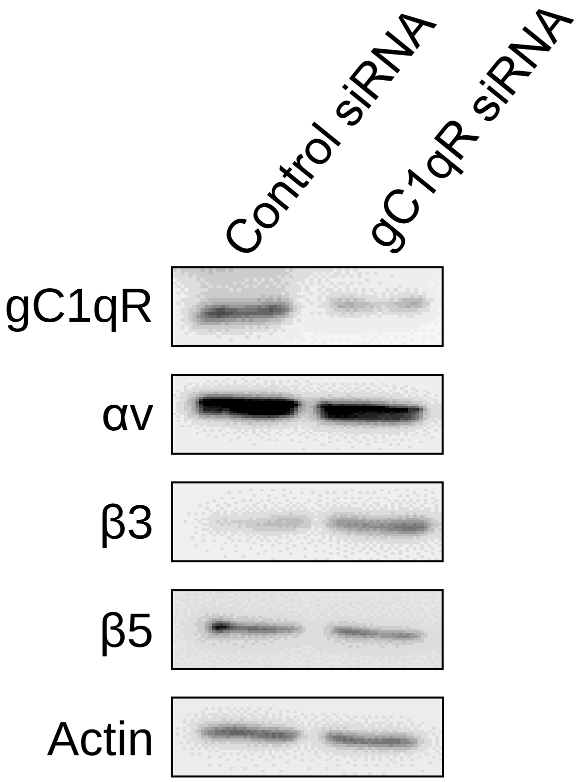


**Fig. S2.** Western blot showing effects of gC1qR siRNA on the levels of the indicated endothelial cell proteins.

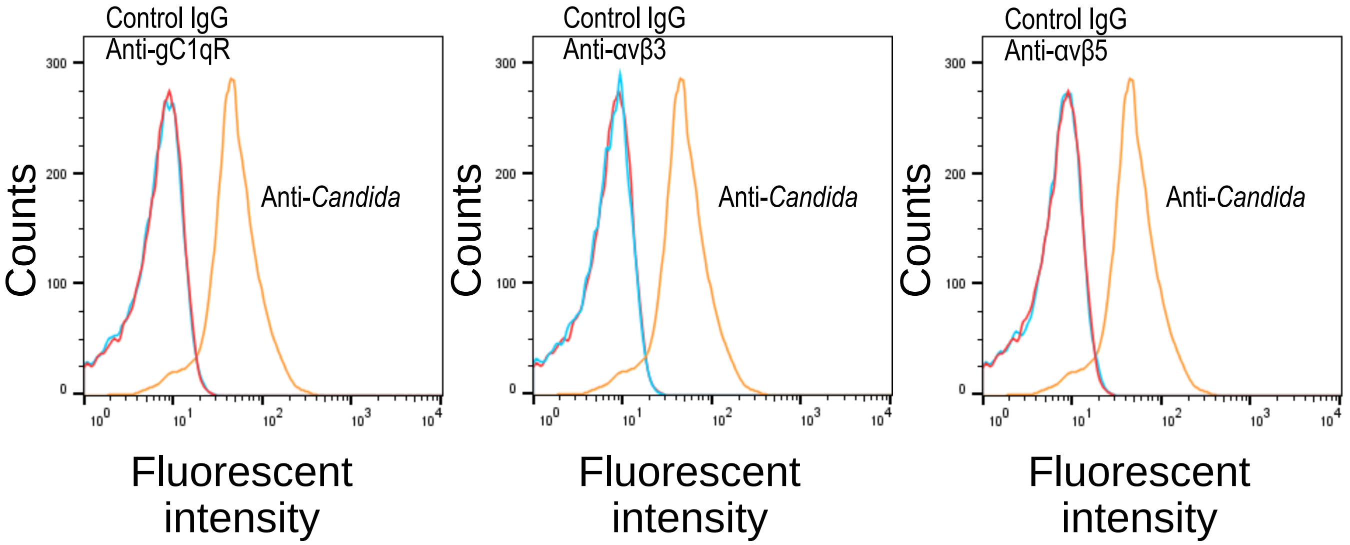


**Fig. S3.** Antibodies against gC1qR, integrin αvβ3, and integrin αvβ5 do not bind to *C. glabrata.* Flow cytometric analysis of *C. glabrata* cells that were incubated with antibodies against gC1qR (clone 74.5.2), integrin αvβ3, and integrin αvβ5. They were also incubated with control IgG and with a polyclonal anti-*Candida* antibody. Each histogram shows the analysis of $10^4$ cells.

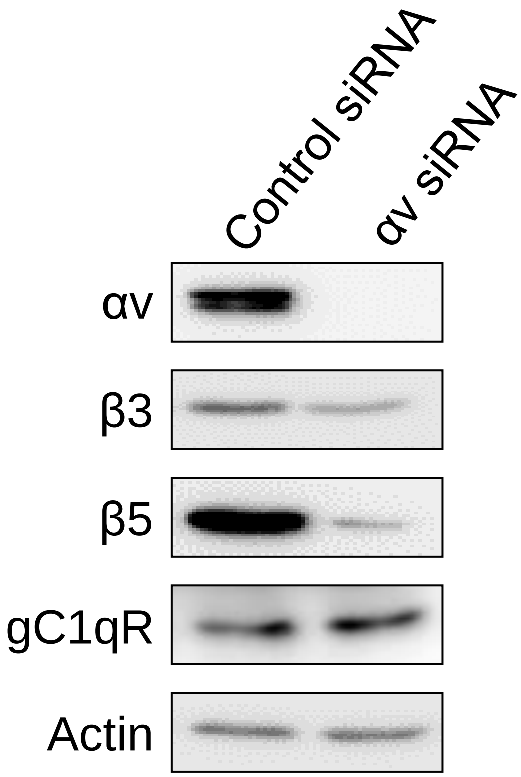


**Fig. S4.** Western blot showing effects of integrin αv siRNA on the levels of the indicated endothelial cell proteins.

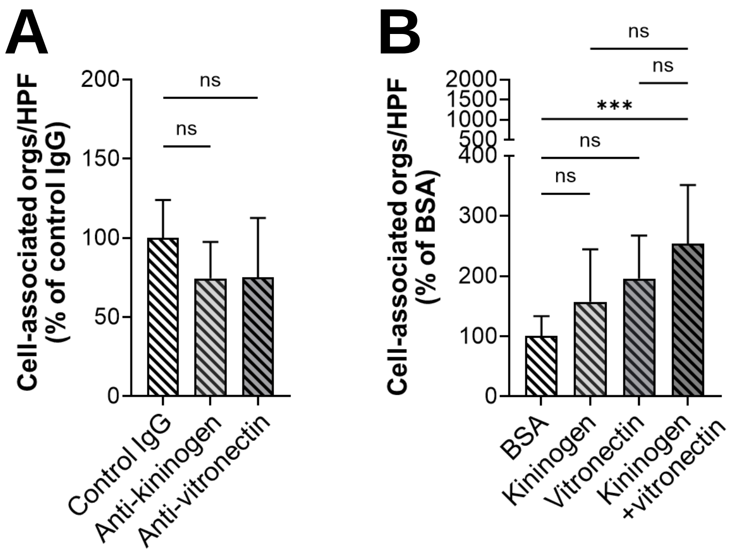


**Fig. S5.** Role of kininogen and vitronectin in mediating the cell-association of *C. glabrata* with human endothelia cells. (A) Effects of antibodies against kininogen and vitronectin on the number of cell-associated organisms. (B) Effects of coating *C. glabrata* with BSA, kininogen, and vitronectin on the number of cell-associated organisms. Data are the mean ± SD of 3 experiments each performed in triplicate. Orgs/HPF, organisms per high power field; ns, not significant; ***$P < 0.001$.

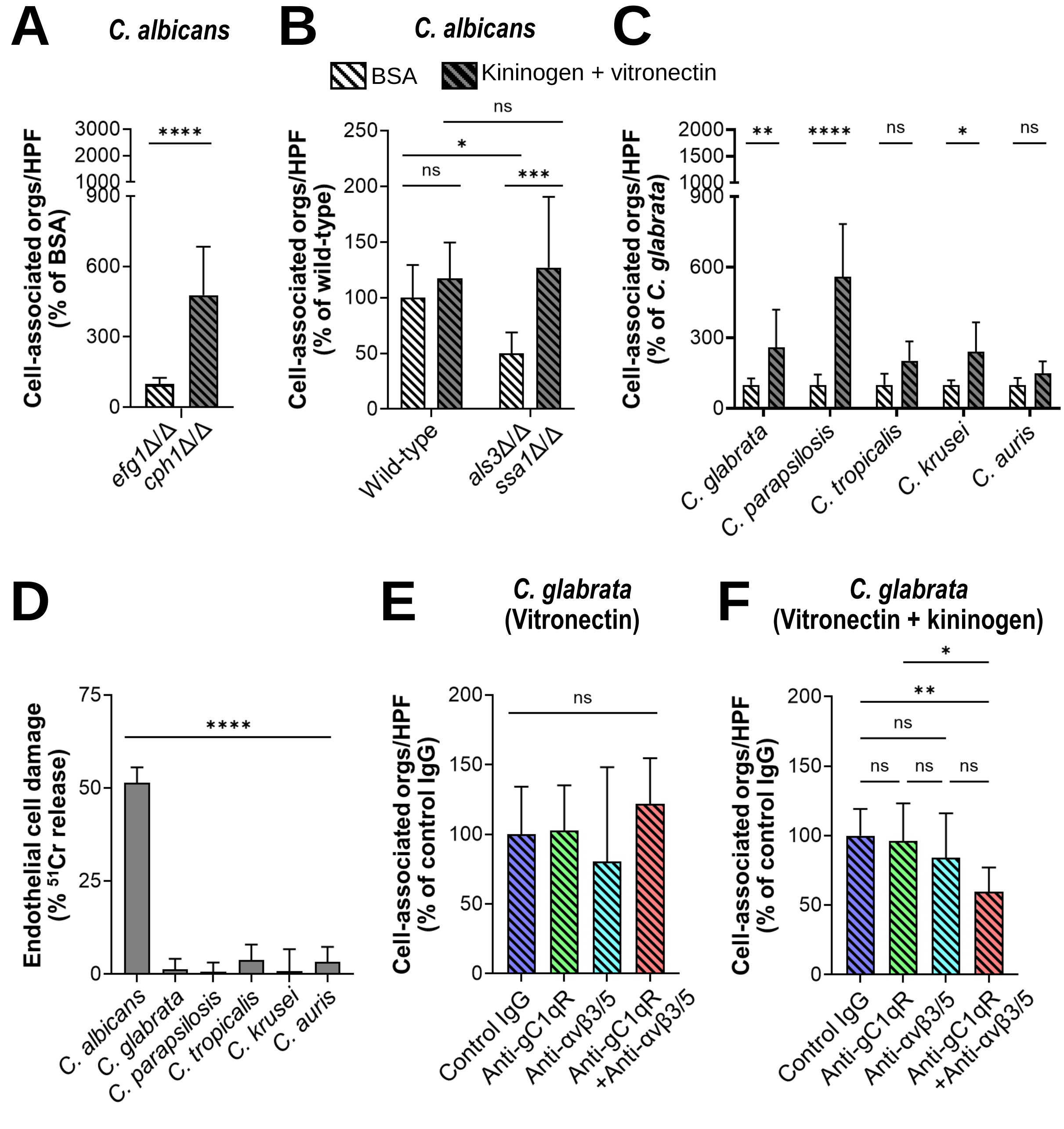


**Fig. S6.** Kininogen and vitronectin interact with gC1qR and αv integrins to induce adherence. (A and B) Effects of BSA or kininogen and vitronectin on the endocytosis the indicated strains of *C. albicans*. (C) Kininogen and vitronectin increase cell-association (adherence) of the indicated *Candida spp.* (D) Endothelial cell damage caused by the cells of the indicated strains that had been coated with kininogen and vitronectin. (E and F) Inhibition of cell-association of *C. glabrata* coated with either vitronectin alone (E) or vitronectin and kininogen (F) by antibodies against gC1qR (clone 74.5.2) and/or integrins αvβ3 and αvβ5. Data are the mean ± SD of 3 experiments each performed in triplicate. Orgs/HPF, organisms per high power field; ns, not significant; *$P < 0.05$, **$P < 0.01$, ***$P < 0.001$, ****$P < 0.0001$.

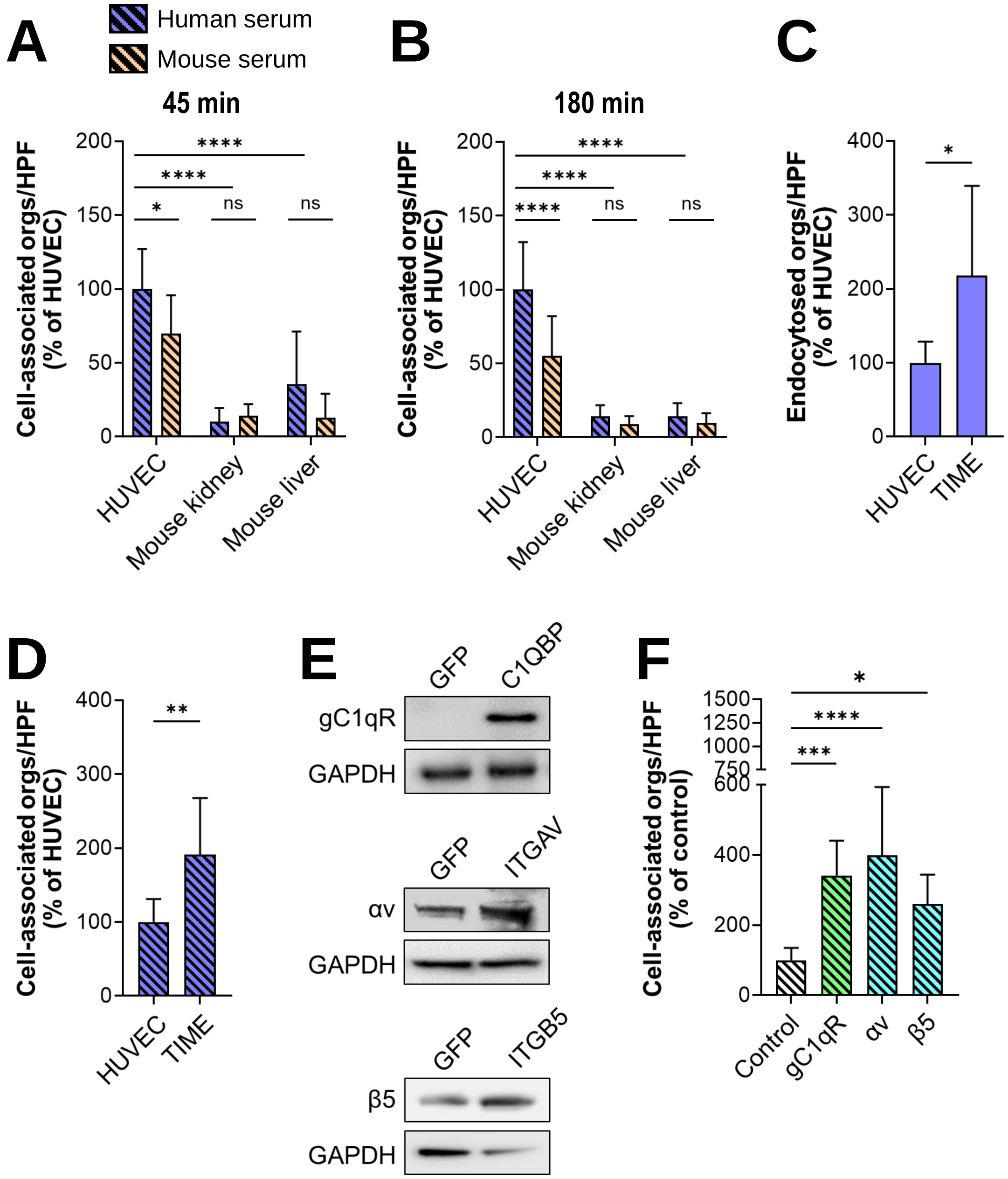


**Fig. S7.** Mouse endothelial cells poorly support bridging molecule-mediated adherence. (A and B) Cell-association of *C. glabrata* coated with either human or mouse serum by the indicated endothelial cells after 45 min (A) and 180 min (B). (C and D) Endocytosis (C) and cell-association (D) of *C. glabrata* coated with fresh human serum by the indicated endothelial cells. (E) Western blot showing the protein levels of gC1qR, integrin αv and integrin β5 in mouse liver endothelial cells transduced with lentivirus containing the indicated human genes. gC1qR was detected with monoclonal antibody 60.1l, which only binds to the human protein. The integrins were detected with antibodies that recognized both human and mouse proteins. (F) Cell-association of *C. glabrata* coated with fresh human serum by mouse liver endothelial cells expressing human gC1qR, integrin αv, or integrin β5. Data in (A-D and F) are the mean ± SD of 3 experiments each performed in triplicate. HUVEC, human umbilical vein endothelial cells; orgs/HPF, organisms per high power field; ns, not significant; TIME, Tert-immortalized microvascular endothelial cells; *$P < 0.05$, **$P < 0.01$, ***$P < 0.001$, ****$P < 0.0001$.
